# Ovarian carcinoma immunoreactive antigen-like protein 2 (OCIAD2) is a novel metazoan specific complex III assembly factor

**DOI:** 10.1101/2020.12.09.417287

**Authors:** Katarzyna Justyna Chojnacka, Karthik Mohanraj, Sylvie Callegari, Ben Hur Marins Mussulini, Praveenraj Elanchelyan, Aleksandra Gosk, Tomasz Banach, Tomasz Góral, Peter Rehling, Remigiusz Adam Serwa, Agnieszka Chacińska

## Abstract

Assembly of the dimeric complex III (CIII_2_) in the mitochondrial inner membrane is an intricate process in which many factors are involved. Despite many studies this process is yet to be completely understood. Here we report the identification of human OCIAD2 (Ovarian Carcinoma Immunoreactive Antigen domain containing protein 2) protein as an assembly factor for CIII_2_. OCIAD2 was found deregulated in several carcinomas and in some neurodegenerative disorders; however its non-pathological role was not elucidated to date. We have shown that OCIAD2 localizes to mitochondria and interacts with electron transport chain (ETC) proteins. Complete loss of OCIAD2 using gene editing in HEK293 cells resulted in abnormal mitochondrial morphology, decrease assembly of both CIII_2_ and supercomplex III_2_+IV and decreased activities of complex I and III. Identification of OCIAD2 as a protein required for assembly of functional CIII_2_ provides a new insight into the biogenesis and architecture of the ETC. Elucidating the mechanism of OCIAD2 action is important both for the understanding of cellular metabolism and for understanding of its role in the malignant transformation.

## INTRODUCTION

Mitochondria are ubiquitous organelles, essential for ATP production by oxidative phosphorylation (OXPHOS). In mammals, OXPHOS relies upon five multisubunit enzyme complexes and two mobile electron carriers embedded within the inner mitochondrial membrane. The first four enzyme complexes (CI-IV) form the electron transport chain (ETC), transfer electrons from electron donors through a series of electrons acceptors, to molecular oxygen, which is coupled to the generation of a proton gradient across the inner membrane that is used by the ATP synthase (CV) to drive ATP synthesis. With exception of the CII, each OXPHOS complex has a dual genetic origin. Subunits encoded by the mitochondrial DNA (mtDNA) are co-translationally inserted into the membrane by matrix-located ribosomes tethered to OXA1 insertase (Hell *et al*., 2001; Haque *et al*., 2010; Ott and Herrmann, 2010). These mitochondrially encoded subunits act as a “seed” around which the rest of the nuclear-encoded subunits are imported from the cytosol through the outer- (TOM) and inner-membrane (TIM) translocase machineries (Neupert and Herrmann, 2007; Chacinska *et al*., 2009; Wiedemann and Pfanner, 2017). In mammalian cells, individual OXPHOS CI, III and IV physically interact to form a variety of supramolecular structures called supercomplexes (SCs I-III_2_, III_2_-IV, I-III_2_-IV_1-4,_ or “respirasome”) (Schafer *et al*., 2006; Althoff *et al*., 2011; Cogliati *et al*., 2016; Gu *et al*., 2016; Letts and Sazanov, 2017). It was proposed that supercomplexes formation stabilizes ETC complexes, facilitates electron transfer, and decreases formation of reactive oxygen species (ROS) but the exact function of supercomplexes is still under debate (Dudkina *et al*., 2010; Milenkovic *et al*., 2017; Lobo-Jarne and Ugalde, 2018) . The assembly of ETC complexes is a convoluted process, due to the nature of the ETC complexes themselves. First, resulted from the dual genetic origin of ETC complexes a tight co-regulation of nuclear and mitochondrial encoded gene expression is required (Mick *et al*., 2012; Rampelt and Pfanner, 2016; Richter-Dennerlein *et al*., 2016; Priesnitz and Becker, 2018). Second, many subunits are embedded in the inner mitochondrial membrane and they are prone to aggregation prior to and during assembly (Chiti *et al*., 2003; Vendruscolo *et al*., 2003; Dobson, 2004). Furthermore, each protein complex contains multiple redox active cofactors that can promote the formation of harmful radicals if not properly assembled to the final complex (Perry *et al*., 2003). As a results a number of proteins so-called assembly factors are needed to coordinate different steps of ETC assembly process (McKenzie and Ryan, 2010).

Assembly factors represent a functionally heterogeneous group of proteins, most of which are specific for the assembly of each ETC complex and they are not part of the final complex. Many assembly factors are conserved between yeast and human, but new factors not related to the yeast proteins are being discovered in human cells (Signes and Fernandez-Vizarra, 2018). Mutations in the assembly factors and subsequent impaired respiratory complexes assembly are associated with a number of pathologies that mainly affect tissues with high energy requirements (Diaz *et al*., 2011; Ghezzi and Zeviani, 2018). Thus, full understanding of the electron transport chain assembly regulation is essential to understand the molecular mechanisms underlying mitochondrial pathology. Despite extensive studies, the exact mechanism by which ETC complexes are assembled is not fully described, and some putative assembly factor proteins remain unknown.

Complex III (or ubiquinol:cytochrome c oxidoreductase), plays a central role in the electron transport chain transferring electrons from coenzyme Q to cytochrome c. All available X-ray diffraction structures (Xia *et al*., 1997; Iwata *et al*., 1998; Zhang *et al*., 1998; Hunte *et al*., 2000) and Cryo-EM structure of CIII (Gu *et al*., 2016) are identical in both shape and subunit composition. CIII is always a symmetrical dimer (CIII_2_), with each monomer being composed of eleven different subunits. Three of them contain the catalytic centers: cytochrome b (MT-CYB), cytochrome c1 (CYC1) and the Rieske protein (UQCRFS1), whereas the other eight subunits (UQCRC1, UQCRC2, UQCRH, UQCRB, UQCRQ, Subunit 9, UQCR10 and UQCR11) do not possess prosthetic groups and do not directly participate in electron transport or proton pumping (Smith *et al*., 2012). The formation of CIII starts with the assembly of two factors UQQC1, UQQC2 to newly synthesized mtDNA-encoded cytochrome *b* (MT-CYTB) (Tucker *et al*., 2013). Binding of the first nuclear-encoded subunits releases these early assembly factors from cytochrome b and the rest of the structural subunits are then incorporated into the nascent complex resulting in the formation of dimeric pre-complex III (Smith *et al*., 2012). Then UQCRFS1 protein is bound with the assistance of 3 assembly factors (BCSL-1, LYRM-7 and TTC-19) (Cruciat *et al*., 1999; Atkinson *et al*., 2011; Ghezzi *et al*., 2011; Sanchez *et al*., 2013; Bottani *et al*., 2017). This step is crucial for CIII_2_ maturation because it turns the enzyme catalytically active. Finally, the last subunit (UQCR11) binds to the CIII_2_, and the assembly is completed. Given the structural similarity between yeast and mammalian CIII as well as identification of several orthologs of yeast assembly factors in humans, the assembly of CIII in mammals is considered to be similar to the one in yeast. However, there is still lack of experimental data regarding some of the assembly steps in the human model. Moreover, assembly factors specific for mammalian CIII such as TTC-19 and BRAWNIN have been also described (Ghezzi *et al*., 2011; Bottani *et al*., 2017; Zhang *et al*., 2020). This make a challenge to unravel unknown mechanisms and factors specific to mammalian CIII biogenesis.

In the present study, we identified and characterized a novel CIII assembly factor specific to metazoan cells, ovarian carcinoma immunoreactive antigen-like protein 2 (OCIAD2). Recent studies explored the origin, evolution and function of OCIAD2 protein that has been shown to be expressed in endosomes and mitochondria (Sinha et al., 2018). In the present study, we explored the functional role of this protein in mitochondria. So far OCIAD2 expression has been implicated in several cancers and neurodegenerative diseases (Han *et al*., 2014; Wu *et al*., 2017; Sinha *et al*., 2018) but its precise function in non-pathological contexts remains unknown. In this report we describe OCIAD2 as a novel CIII assembly factor of the mitochondrial respiratory chain in human cells.

## RESULTS

Sequence analysis revealed that OCIAD2 is conserved in metazoan (Supplementary Figure 1) and no yeast homolog could be identified. Moreover, OCIAD2 amino-acid sequence is devoid of canonical cleavable N-terminal mitochondrial targeted presequence and indicate the presence of 2 α-helical stretches of high hydrophobicity that are probably transmembrane (TM) domains (Figure 1A). To check whether OCIAD2 despite the lack of predicted cleavable presequence is mitochondrial localized we analyzed its cellular localization by immunofluorescence microscopy using an antibody specifically directed against native OCIAD2. OCIAD2 signal largely colocalized with Mitotracker Orange™ (Figure 1B). We additionally prove the mitochondrial localization by performing cellular fractionation, where native OCIAD2 was predominantly localized to mitochondria (Figure 1C, lane 3). The association of OCIAD2 with the mitochondrial membranes was analyzed by treatment of mitochondria with Na_2_CO_3_ which causes the release of peripheral membrane associated proteins at pH 10.8. OCIAD2 was resistant to carbonate extraction (Figure 1D, line 3), alike the membrane integrated proteins TOMM20, MIC60 and TIMM29, indicating that it is very likely an integral membrane protein. Next, we examined the submitochondrial localization of OCIAD2 by performing a hypo-osmotic swelling (Figure 1E). In intact mitochondria, native OCIAD2 was protected from the degradation by the outer membrane (OM), similar to other proteins localized inside mitochondria, such as COA7 (intermembrane space, IMS), TIMM22, TIMM29 (inner membrane, IM), and HSP60 (matrix) (Figure 1E, lane1). As expected, the OM protein TOMM20 was efficiently degraded by proteinase K in intact mitochondria (Figure 1E, lane 2). Upon disruption of the OM by hypo-osmotic swelling, the IMS becomes accessible to protease treatment. Under these conditions using antibody against N-terminal part of OCIAD2 we observed digestion of OCIAD2 by the protease similarly to IMS and IM proteins (Figure 1E, compare lines 2 and 4). Consistently, we concluded that OCIAD2 is an inner membrane protein with its N-terminal part exposed to the IMS.

**Figure 1:**
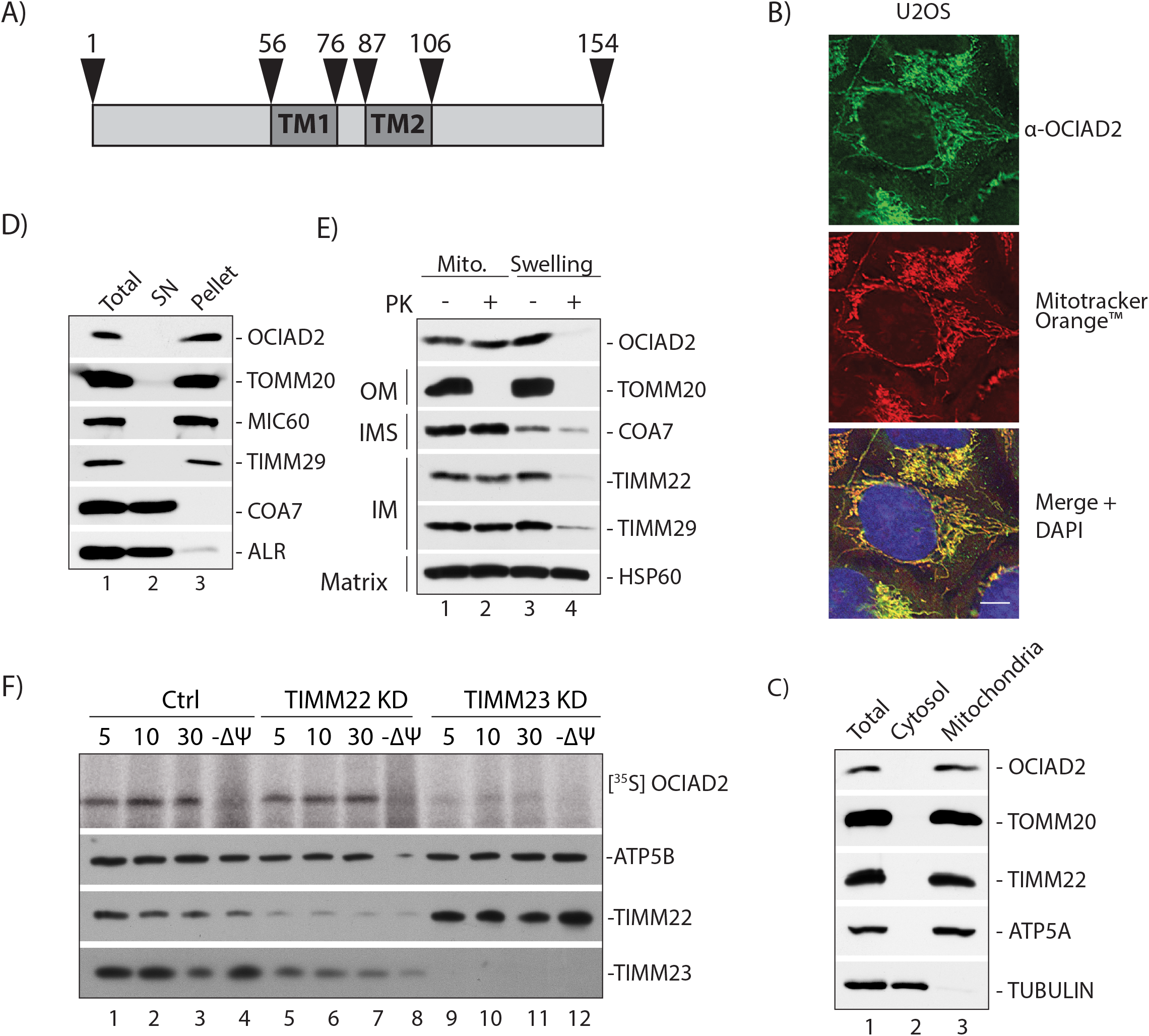
OCIAD2 is an inner mitochondrial membrane protein. (A) Primary sequence of OCIAD2. Dark grey boxes area depicts the predicted transmembrane segments. (B) U2OS were immunolabeled using antibody against OCIAD2 and stained with Mitotracker Orange™ to visualize the mitochondria. Scale bar, 8 µm. (C) Subcellular fractionation of HEK293 cells. (D) Carbonate extraction of KHE293 calls at Ph 10.8. After differential centrifugation, samples were analyzed by SDS/PAGE and western blotting. (E) Isolated mitochondria from HEK293 cells were subjected to submitochondrial fractionation. Mito chondria (lanes 1 and 2); hypotonic swelled mitochondria (lanes 3 and 4) were treated with (+) or without (−) Proteinase K (PK). (F) Radiolabeled OCIAD2 precursors were imported into mitochondria isolated from Control and TIMM22KD or TIMM23KD cells for indicated times in the presence or absence of a membrane potential (Ψ) across the IM and treated with proteinase K (PK; 50 mg/mL). Samples were analyzed by SDS-PAGE followed by autoradiography or immunoblotting using ATP5B, TIMM22 or TIMM23-specific antibodies.

The majority of inner membrane protein precursors are synthesized in the cytosol and imported into the mitochondria by TIM22 or TIM23 translocases. TIM22 is responsible for the import of the precursor proteins carrying internal targeting signals whereas TIM23 facilitates the import of the presequence-containing preproteins (Neupert and Herrmann, 2007; Chacinska *et al*., 2009). It should be also noted that many proteins from the inner membrane does not resemble typical substrates for TIM22 or TIM23 translocase (Reinhold *et al*., 2012; Turakhiya *et al*., 2016; Rampelt *et al*., 2020). To distinguish import pathway that OCIAD2 uses for the integration into the inner membrane we apply an *in vitro* import assay. In brief, radioactive [^35^S]-OCIAD2 was synthesized in rabbit reticulocyte lysate and imported into mitochondria isolated from control (Ctrl) HEK293 cells and into HEK293 cells depleted of TIM22 or TIM23 translocases by siRNA (Figure 1F). In case of the ctrl cells (Figure 1F, lines 1-4) and TIMM22 knock-downed cells (Figure 1F, lines 5-8) OCIAD2 was imported into a protease-protected location, in a time-dependent manner. In contrast, OCIAD2 import into the mitochondria lacking TIMM23 was inhibited (Figure 1F, lines 9-12). This indicates that despite the absence of a presequence, OCIAD2 requires TIM23 for its import into the mitochondria.

### OCIAD2 interacts with respiratory chain complexes

In order to functionally characterize OCIAD2 in the human mitochondria we catalogue OCIAD2 protein binding partners by performing affinity purification of ^FLAG^OCIAD2 under non-denaturing conditions followed by quantitative mass spectrometry. We first generated stable tetracycline-inducible Flp-In™ T-REx™ 293 cell line expressing OCIAD2 protein with a N-terminal FLAG-tag. Confocal microscopy showed co-localization of FLAG-OCIAD2 with the Mitotracker™ Orange, thus confirming mitochondrial localization of the fusion protein (Figure 2A). ^FLAG^OCIAD2-expressing cell line and the control cells were then differentially labeled via stable isotope labeling by amino acids (SILAC) and mixed early in the protocol, in order to minimize potential experimental errors resulting from parallel mitochondria isolation and subsequent processing steps. Following FLAG-based enrichment, proteins were digested with trypsin and analyzed by shotgun proteomics. Next, we filtered the list of identified proteins by their annotation in the Integrated Mitochondrial Protein Index (IMPI) reference set of genes deposited in the MitoMiner database ver. 4.0 (Smith and Robinson, 2019). Amongst 180 mitochondrial proteins identified in our analysis, we assigned 48 as putative binding partners of OCIAD2, based on the mean fold enrichment value >2 in the comparison of intensities of proteins derived from ^FLAG^OCIAD2 vs control cells. The list of identified mitochondrial proteins (Table S1) was then subjected to gene ontology (GO) annotation, and to a term enrichment analysis (Supplementary Figure 2). One of the most enriched GO terms was related to the respiratory electron transport chain (ETC). Indeed, a large pool (29) of proteins constituting ETC CI, III and IV co-purified with ^FLAG^OCIAD2 (Figure 2B). In contrast, some large groups of proteins, sharing common terms, e.g. ATP synthase (9), mitoribosomal protein (37) were not bound specifically to the ^FLAG^OCIAD2 as judged by the apparent lack of enrichment (fold enrichment <2) (Figure 2B). Next, we validated the proteomic results, by Western-Blot analysis employing antibodies specific towards selected proteins identified in the mass spectrometric analysis (Figure 2C). Indeed, with this technique, we confirmed that proteins of ETC complexes I, III and IV were efficiently co-purified with the bait, in contrast to mitoribosomal proteins or ATP synthase (Figure 2C).

**Figure 2:**
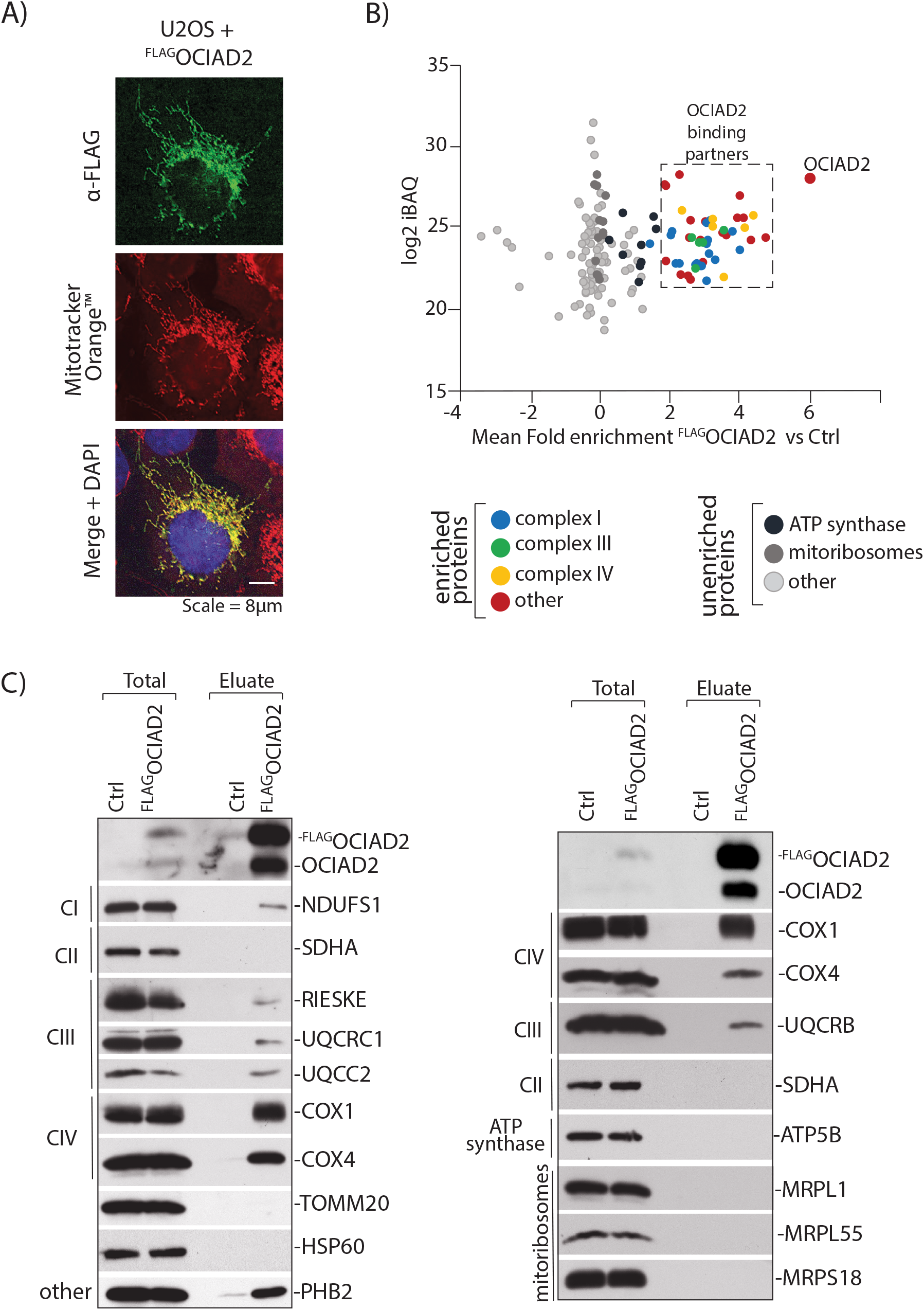
OCIAD2 interacts with proteins from electron transport chain. (A) U2OS cells expressing ^FLAG^OCIAD2 were immunolabeled using antibodies against FLAG. Mitotracker Orange™ was used to visualize the mitochondria. Scale bar, 8 µm (B) Mitochondria isolated from control HEK293 Flp-in WT cells (SILAC light) and from the cells expressing ^FLAG^OCIAD2 protein (SILAC heavy) were mixed and solubilized in digitonin-containing buffer following a FLAG-targeting immunoprecipitation. Fold enrichment of proteins co-immunoprecipitated with ^FLAG^OCIAD2 vs Ctrl (x-axis) was correlated with intensity-based absolute quantification (iBAQ) values, which is a measure of protein abundance (y-axis) categories. For the entire mitochondrial protein list, see Table 1. (C) Control mitochondria and mitochondria expressing ^FLAG^OCIAD2 were solubilized in digitonin-containing buffer and subjected for immunoprecipitation with FLAG affinity resin. Samples were analyzed using SDS-PAGE and immunoblotting with the indicated antibodies. Total 1%; Eluate 100%

### Loss of OCIAD2 causes mitochondria morphology and growth defect and influences mitochondrial bioenergetics

The observation that OCIAD2 binds to many proteins from ETC complexes led us to investigate if OCIAD2 has an influence on the activity of these complexes. To this end, we generated a knockout of OCIAD2 (OCIAD2-KO) using CRISPR/Cas9-mediated disruption of both the alleles in HEK293 cells. The second exon of the gene was targeted and the selected knockout clone was a compound homozygote for a 66-nucleotide deletion (1_66del) encoding the N-terminal part of OCIAD2 protein (Figure 3A). OCIAD2-KO cell line was verified by sequencing of the genomic locus (Figure 3A) and Western blot analysis (Figure 3B). Reduction of OCIAD2 levels resulted in a growth phenotype (Figure 3C), and abnormal mitochondria morphology compared to WT cells (Figure 3D) which suggests alterations in mitochondrial functions. Hence we aimed to analyze in detail the performance of mitochondria in the OCIAD2-KO cells. To that end we analyzed the mitochondrial respiration by measuring the oxygen consumption rates (OCRs) in wild type (WT) and OCIAD2-KO cells. First, we used intact cells to keep undisturbed cellular environment and to minimize potential artefacts due to mitochondrial preparation (Figure 4 A-B). Under these conditions, deletion of OCIAD2 had no significant impact on basal respiration compared to WT cells. However, the maximal respiration induced by the protonophore CCCP was significantly lower in OCIAD2-KO cells compared to WT (66% residual respiration, *p*<0.05) (Figure 4B). The reduction of maximal respiration led us to assess CI, CIII and IV activities coupling to oxygen consumption as the final electron acceptor. Mitochondria depleted of OCIAD2 exhibited decreased oxygen consumption driven by pyruvate and malate (CI substrates electron donors - Figure 4C), and by glycerol-3-phosphate (CIII substrate electron donor - Figure 4D). Complex IV activity was not changed upon OCIAD2 deletion (Figure 4E). Altogether, OCIAD2 does not affect the basal activity of mitochondria neither oxygen consumption couple to ATP production indicating no influence on mitochondria activity under metabolic non-stress situation. Nevertheless, reduction in maximal respiration and spare capacity coupled to a reduction in CI and CIII activities indicates that OCIAD2 depleted cells are exposed to metabolic stress. These data confirm that OCIAD2 is essential for cellular bioenergetics, potentially through regulation of the electron transport chain.

**Figure 3:**
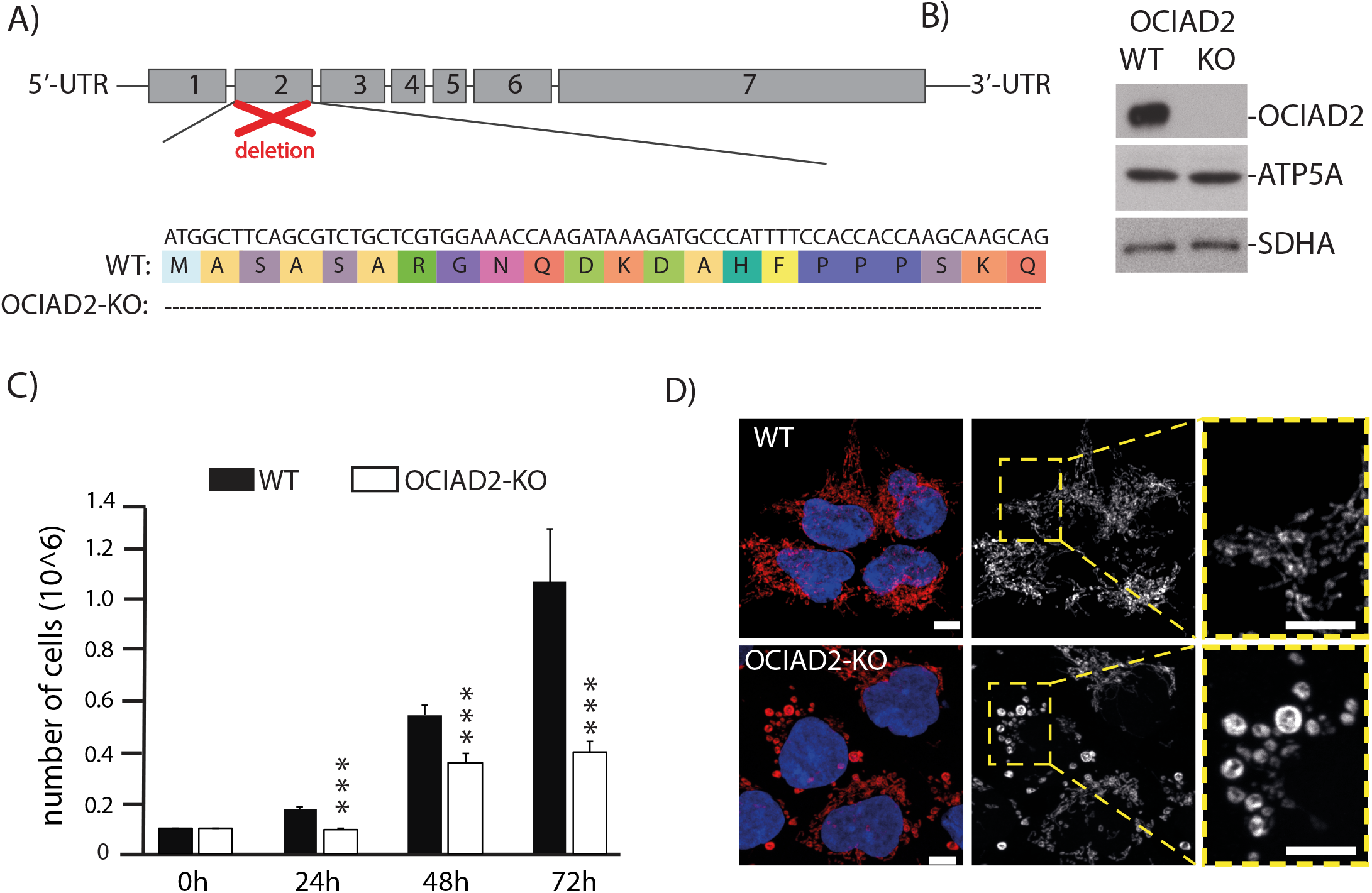
OCIAD2-KO cells display abnormal mitochondria morphology and proliferation. (A) Knockout cell line of OCIAD2 (OCIAD2-KO) was created in HEK293 cells by using CRISPR/Cas9 technology. Alignment of respective mutated region compared to WT shows deletion of 66 aa from the N-terminal part of protein. (B) Western blot results confirming OCIAD2 deletion (C) WT and OCIAD2-KO cells were seeded 24h prior experiment at concentration of 0.1 × 10 /well in the growth medium. After 24h, 48h and 72h of culture at 37°C with 5% CO2, cells were harvested by trypsinization and counted. Data are shown as the mean ± SD (n = 5) ***p<0.01(two-tailed Student’s t -test). (D) WT and OCIAD2-KO cells were immunolabeled using antibody against cyclophilinD to visualize the mitochondria. Scale bar, 10 µm.

**Figure 4:**
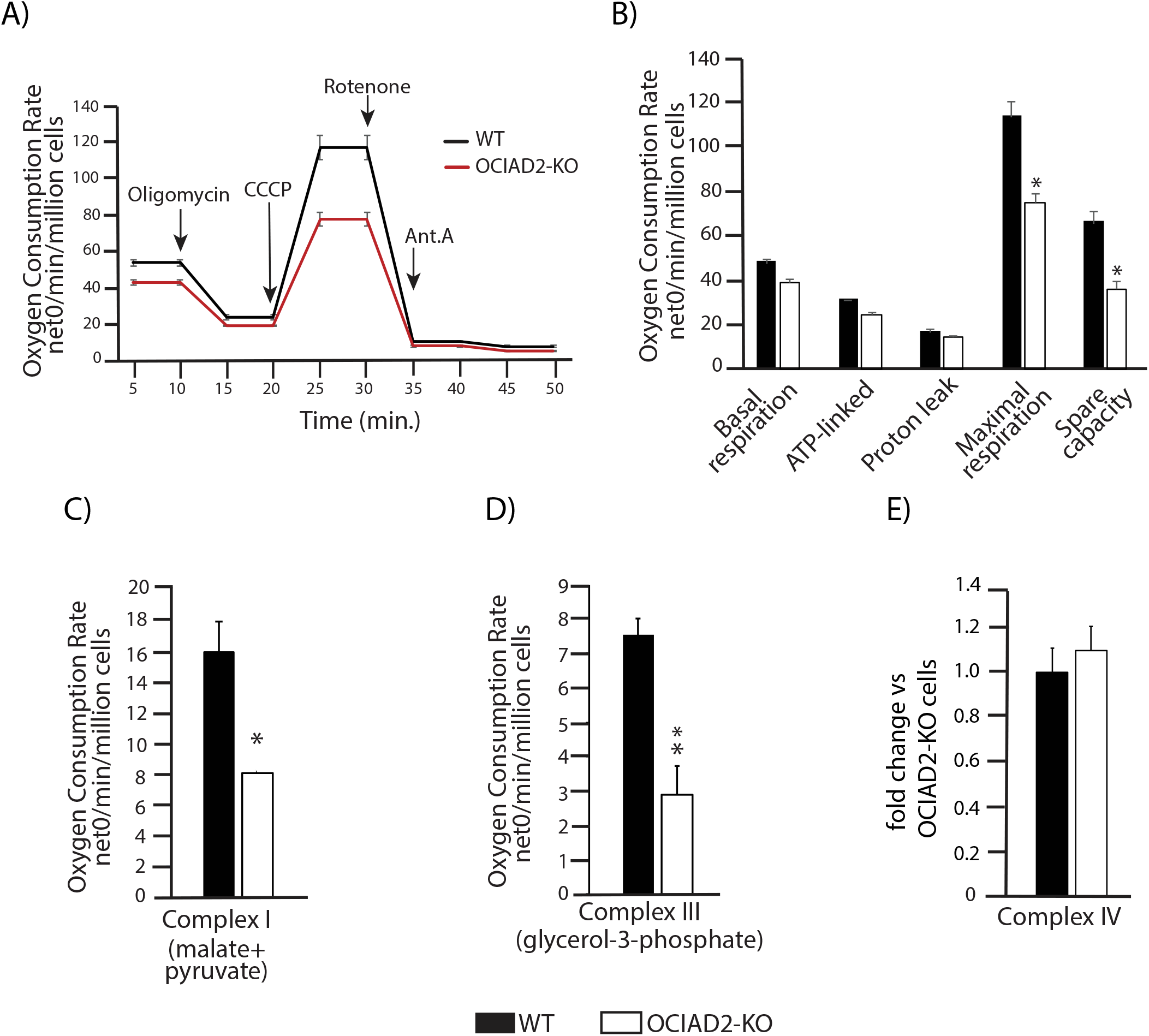
OCIAD2 influences bioenergetic in mitochondria. (A) Oxygen consumption rates (OCR) in intact OCIAD2-KO cells cultured in glucose. Oligomycin (ATP synthase blocker), CCCP (uncoupler), rotenone (CI inhibitor) and antimycin A (CIII inhibitor), were added at indicated time points. OCR was normalized to number of cells. Data are presented as the mean ± SD (n = 3). *p < 0.05. (two-tailed Student’s t -test).(B) Basal, ATP-linked, proton leak, maximal OCR, and spare respiratory capacity were quantified from the original data presented in A. Data are shown as the mean ±SD (n=3).*p < 0.05, **p<0.01 (two-tailed Student’s t -test).(C) Respiration of digitonin-permeabilized HEK293 cells assessed in the presence of Pyr-Mal: pyruvate (5 mM) and malate (0.5 mM) to evaluate CI activity (D) Respiration of digitonin-permeabilized HEK293 cells assessed in the presence of glycerol-3-phosphate (5 mM) to evaluate CIII activity. (E) CIV activity was assessed by oxidation of cytochrome c in digitonized cellular extracts from the OCIAD2-KO and WT cells. Data are shown as the mean ± SD (n = 5)(two-tailed Student’s t -test).

### OCIAD2 is required for the assembly of respiratory chain complexes

As OCIAD2 has an impact on ETC complexes activity we further analyzed their assembly by Blue Native-PAGE (BN-PAGE). Mitochondria from OCIAD2-KO cells were solubilized in digitonin or in n-Dodecyl β-D-maltoside (DDM) containing buffer which are frequently used detergents in the isolation of membrane protein complexes prior to analysis by BN-PAGE and immunodetection (Figure 5A). We used both buffers since digitonin better preserves native supercomplexes, whereas the harsher DDM induces supercomplexes separation into subcomplexes (Reisinger and Eichacker, 2008). These experiments revealed that fully assembled CIII_2_ as well as III_2_+IV-containing supercomplexes were downregulated in the OCIAD2-KO cell line, as shown in digitonin and DDM treated mitochondria (Figure 5A). We also did not observed changes in the bands pattern corresponding to SC I+III_2_+IV, so called respirasome (Figure 5A). The fully assembled CIV and CV were not affected by the OCIAD2 deletion as shown in DDM treated mitochondria (Figure 5A). In order to check if changes in the assembly of ETC complexes are results of decrease in steady state levels of proteins constituting these complexes we performed Western blot analysis using specific antibodies against selected ETC proteins (Figure 5B). However, these analyses shown no mayor impact of OCIAD2 deletion on steady state levels of ETC proteins. These results suggest that that OCIAD2 is needed for structural rearrangements of ETC complexes but does not participate in ETC protein homeostasis. To validate the phenotype of OCIAD2-KO cells we performed a transient knockdown of OCIAD2 gene using siRNA. Importantly, the alterations in the ETC complexes assembly and their steady state levels in OCIAD2 siRNA mediated knock-down cells were identical to the one observed in the CRISPR/Cas9 OCIAD2-KO cell line (Figure 5C-D), thus validating the causative gene for this phenotype. With the aim of better understand molecular changes induced in mitochondria upon deletion of OCIAD2 we performed a comparative label-free quantitative proteomic analysis of protein levels in the mitochondria isolated from OCIAD2-KO and WT HEK293 cells (Figure 4E). Among all identified 998 mitochondrial proteins (Table S2) we did not observe changes in the steady state levels of proteins constituting CI, CIII and CIV which is in the line with our Western blot results (Figure 5B).

**Figure 5:**
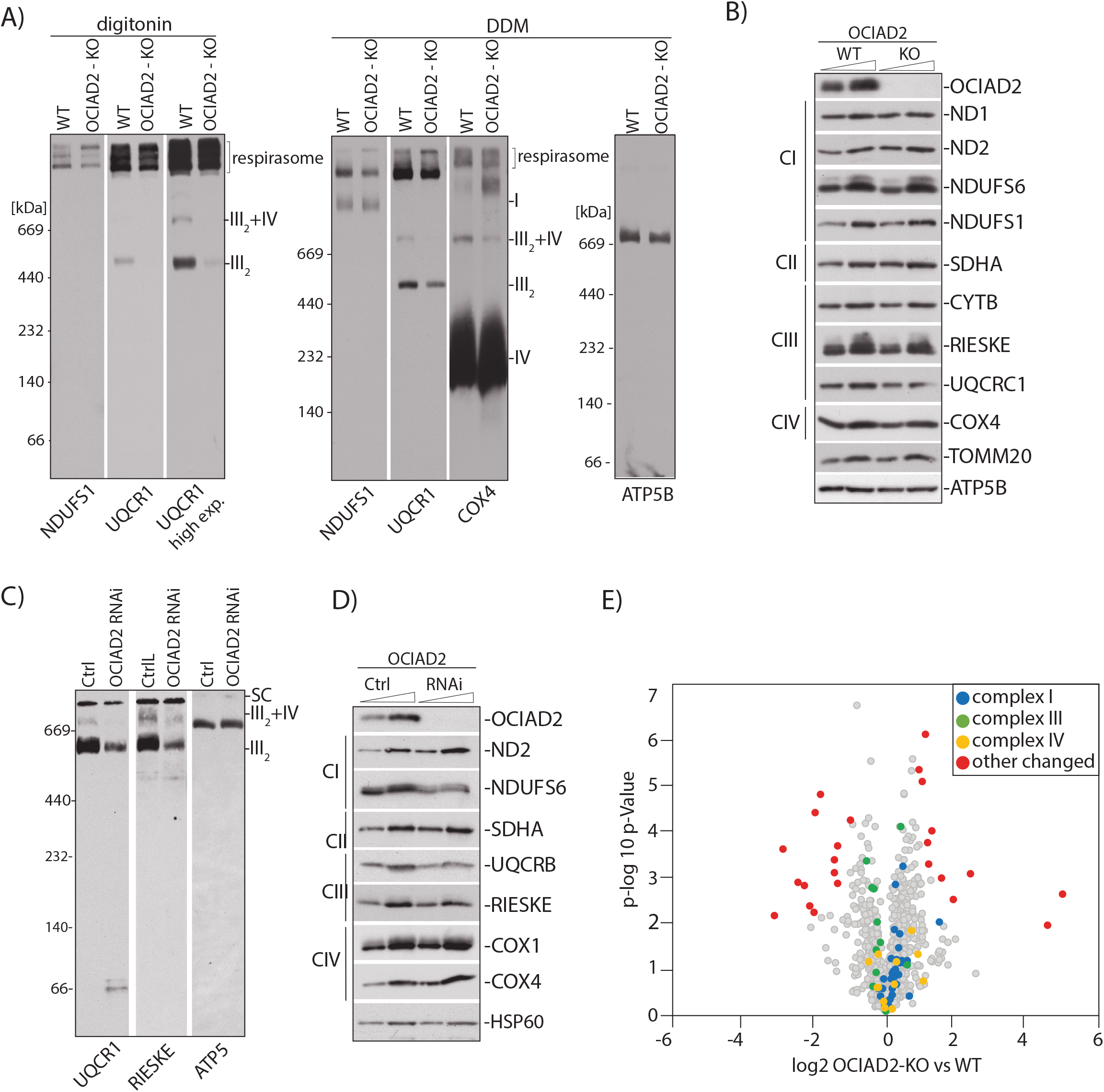
OCIAD2 is needed for the proper assembly of CIII_2_ and SC CIII_2_+IV. (A) Isolated WT or OCIAD2-KO mitochondria were solubilized in 1% digitonin or 0.5 % DDM and resolved using BN-PAGE, followed by western blotting and detection using the indicated antibodies. (B) WT and OCIAD2-KO mitochondria were analyzed by western blotting. (C) Isolated WT or OCIAD2 siRNA mitochondria were solubilized in 1% digitonin, resolved by BN-PAGE, followed by western blotting and immunodetection. (D) Isolated WT and OCIAD2 siRNA mitochondria were analyzed by western blotting. (E) Volcano plot representation of proteins changed upon OCIAD2 deletion. Mitochondria isolated from WT and OCIAD2-KO were lysed in 0.1%TFA, desalted and analyzed by LC-MS/MS (n = 3). Blue, green and yellow color indicate proteins belonging to complex I, III_2_ and IV of ETC, respectively.

The identification of large pool of CI, CIII and CIV proteins as binding partners of OCIAD2 led us to check whether OCIAD2 forms a common complex with ETC subunits. We performed immunopurification of ^FLAG^OCIAD2 and resolved the eluate fraction by BN-PAGE. Using antibody against Rieske and UQCRC1 from complex III we detected OCIAD2 interaction with CIII_2_, SC III_2_+IV and with respirasome (I+III_2_+IV). We also detected interaction of OCIAD2 with respirasome using antibody against CI - NDUFS1. (Figure 6A). We then concluded that OCIAD2 acts on both: dimeric complex III and on respirasome. This prompted us to distinguish whether OCIAD2 is a factor needed for the assembly of particular complex (I or III) or it is dispensable for supercomplexes formation. We focused on CI and CIII since we observed alteration in their composition and activities while no alteration in the case of CIV (Figure 5A and 4C-E). Thus, we assessed import and assembly of the radiolabeled UQCRB into isolated WT and OCIAD2-KO mitochondria by BN-PAGE analyses. As a control we used mitochondria with dissipated membrane potential to ensure that the protein import and assembly depends on the membrane potential across the inner mitochondrial membrane. CIII subunit UQCRB was incorporated into CIII_2_-containing structures in the WT mitochondria, whereas in OCIAD2KO mitochondria incorporation of UQCRB into CIII_2_ was not observed.

**Figure 6:**
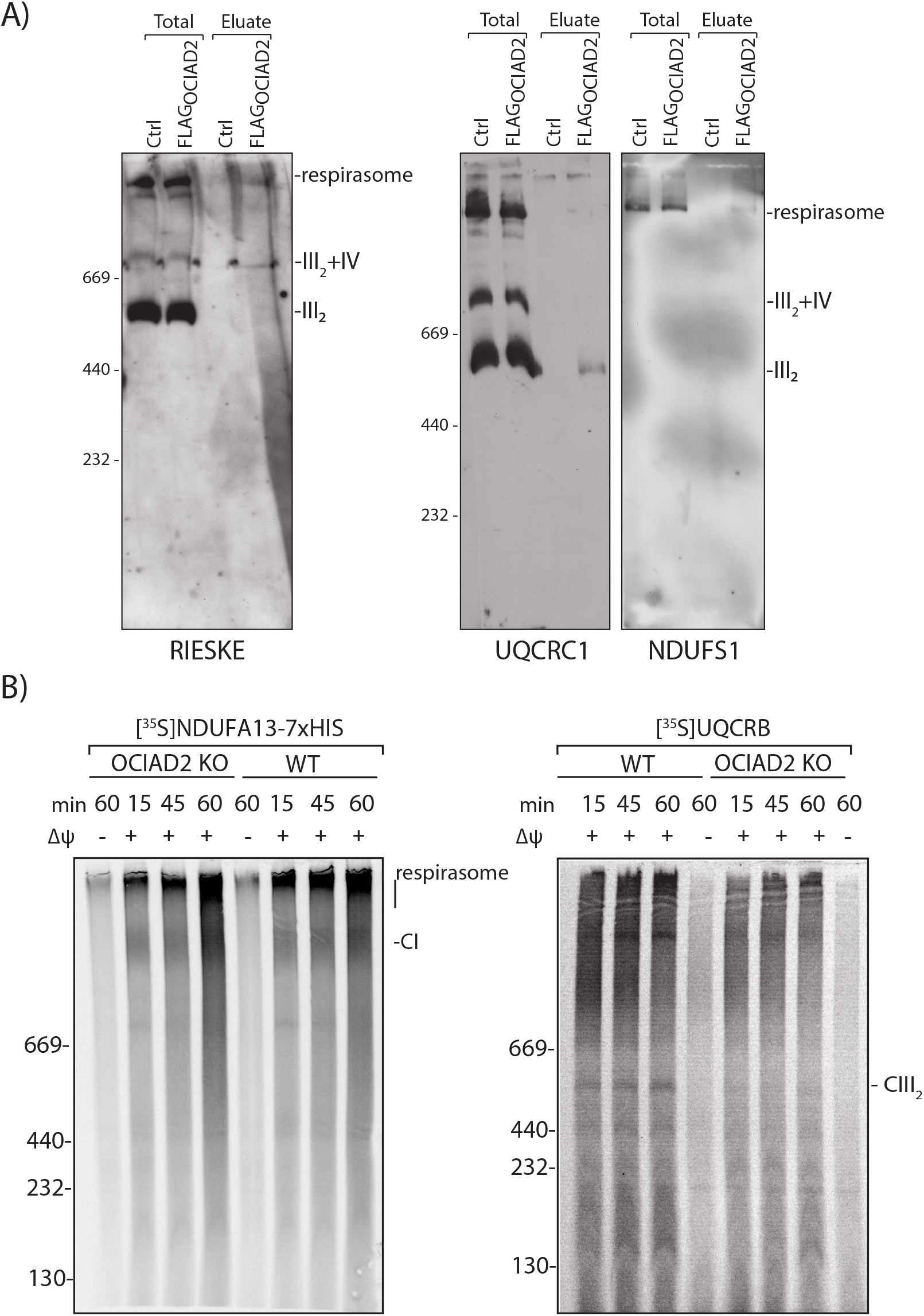
OCIAD2 is involved in the import and assembly of CIII_2_. (A) ^FLAG^OCIAD2 was immunopurified and eluate was resolved by BN-PAGE. (B) Radiolabeled UQCRB, NDUFA13 were imported into isolated mitochondria from wild-type HEK293 and OCIAD2-KO cells, for the indicated times, in the presence or absence of membrane potential (Δψ) at 37°C. Samples were analyzed by BN-PAGE and digital autoradiography.

## DISCUSSION

The CIII, or the cytochrome bc_1_ complex, is structurally similar in yeast and mammals and the current knowledge about CIII assembly factors was obtained mostly by extrapolating the function of well-studied yeast orthologues to mammalian systems. However, some factors specific to higher Eukaryotes have been discovered. Two mammalian assembly factors known to play a role in CIII_2_ biogenesis are: TTC19 and BRAWNIN. The former was discovered in samples associated with a CIII-defective, progressive mitochondrial encephalopathy (Ghezzi 2011), and the latter was found during screening of the small open reading frame (sORF)-encoded peptides (SEPs) in human peptidome (Zhang *et al*., 2020). In this report we describe a new molecular role of the metazoan specific protein, OCIAD2 in the stabilization and assembly of CIII in human cells.

OCIAD2 was originally discovered as an immunoreactive antigen in patients with ovarian cancer (Luo *et al*., 2001). Later OCIAD2 was reported dysregulated in various types of cancer, however, its regulatory roles in cancer progression is still unclear and appears to be tumor type-dependent (Nagata *et al*., 2012; Wu *et al*., 2017; Sakashita *et al*., 2018). In addition OCIAD2 was also implicated in Alzheimer disease (Han *et al*., 2014). OCIAD2 has been previously shown to localize predominantly to the mitochondria (Sinha *et al*., 2018), but its physiological role in the non-pathological context in this organelle remains obscure.

In this work we used a combination of OCIAD2 immunodetection, mass spectrometry-based proteomics, and mitochondrial *in vitro* import assays to demonstrate that OCIAD2 associates with different CIII-containing structures, including CIII_2_, SC III_2_+IV and SC I+III_2_+IV (respirasome). The absence of OCIAD2 causes decrease in the level of assembled CIII_2_ and the SC III_2_+IV with no changes in respirasome formation. This is in accordance with previous observations suggesting that the biogenesis of the SC III_2_+IV is not necessary for respirasome formation (Perez-Perez *et al*., 2016). The fact that respirasomes (but not SC III_2_+IV) are present in OCIAD2-KO mitochondria provides support for a role of OCIAD2 in the stabilization of SC III_2_+IV. These findings also suggest the existence of independent regulatory mechanisms for the biogenesis and turnover of SC III_2_+IV and the respirasomes. Possibly, there is a specific, yet to be-understood, physiological importance of SC III_2_+IV. We also observed reduction in the activities of CI and CIII which is in line with the well-established correlation between severe CIII deficiency and CI impairment (Lamantea *et al*., 2002; Acin-Perez *et al*., 2004; Barel *et al*., 2008; Tucker *et al*., 2013; Wanschers *et al*., 2014; Feichtinger *et al*., 2017; Protasoni *et al*., 2020). Furthermore, OCIAD2KO cells exhibit proper formation and assembly of individual CI and CIV which may result from the fact that respirasome is also not affected in OCIAD2-KO cells. This is in agreement with the study by Protasoni et al. (2020) in which association of respirasome formation was shown to proceed the maturation of stable individual CI and CIV (Protasoni *et al*., 2020). We also show that the deletion of OCIAD2 does not affect steady-state levels of protein components of CIII, but rather the amount of fully assembled CIII_2_. In fact, OCIAD2 deletion does not affect steady-state levels of all detected OXHPOS proteins. Since CIII2 is the necessary component of respirasome, and the formation of respirasome was not affected by OCIAD2 deletion, it is conceivable that this protein may act as a factor stabilizing matured CIII_2_ rather than a factor needed for its maturation. This would make OCIAD2 function similar to that of COX7A2L or UQCC3A, both interacting with CIII_2_ and stabilizing SC III_2_+IV (Desmurs *et al*., 2015; Perez-Perez *et al*., 2016). The similarities between these proteins may suggest their cooperation in the function however this requires further studies to prove such functional interactions. The specific mechanism by which OCIAD2 promotes CIII assembly and/or stability requires further investigation.

We also report here abnormal mitochondrial morphology in cells depleted of OCIAD2, which is not surprising since mitochondrial morphology has been reported to be tightly connected with mitochondrial function. While we cannot entirely exclude that OCIAD2 has some direct role in maintaining the morphology of mitochondria, our results suggest a simpler explanation that altered respiration of OCIAD2-KO mitochondria is the cause of morphological changes. These results are in line with the results of previous studies, which showed a change in mitochondrial morphology in patients with CIII deficiency (Slipetz *et al*., 1991).

Up to now there is no data on patients with mitochondrial dysfunction resulting from mutations in OCIAD2. Nevertheless, identification of OCIAD2 as a assembly/stabilizing factor for CIII_2_ and SC III_2_+IV opens up the possibility of screening for the presence of pathogenic mutations in the OCIAD2 gene in as yet unresolved cases of mitochondrial disease associated with CIII_2_ deficiency. Additionally, since OCIAD2 has been implicated in various cancer types with no common mechanism reported, and mitochondrial dysfunction is a hallmark of cancer, it can’t be excluded that dysfunctional mitochondria due to OCIAD2 loss lead to the cellular changes resulting in cancer. However, the causal relationship between OCIAD2 function in the mitochondrial CIII_2_ and cancer, requires further deep-gain investigation.

## MATERIAL AND METHODS

### Cell lines and growth conditions

For this studies we used Human embryonic kidney cell lines (HEK293T-Flp-InTM T-RExTM) or HEK293 cells purchased from American Type Cell Collection (ATCC; www.atcc.org).Cells were cultured at 37 °C in a 5% CO_2_ atmosphere. The cell lines were sustained in Dulbecco’s modified Eagle’s medium (DMEM) with high glucose content (4500 mg/L), supplemented with 10% (v/v) fetal bovine serum (FBS), 2 mM L-glutamine and 1% (v/v) penicillin-streptomycin. For SILAC analysis, cells were grown for 5 passages on DMEM medium lacking arginine and lysine, supplemented with 10% (v/v) dialyzed fetal bovine serum, 600 mg/L proline, 42 mg/L arginine hydrochloride (^13^C_6_, ^15^N_4_-arginine in ‘heavy’ media), and 146 mg/L lysine hydrochloride (^13^C_6_, ^15^N_2_-lysine in ‘heavy’ media) (Cambridge Isotope Laboratories, Tewksbury, MA, USA). Stable tetracycline-inducible cell line (^FLAG^OCIAD2) was generated using the Flp-IN T-REx System (Thermo Fisher Scientific). Briefly, Flp-IN T-REx HEK293T cells were co-transfected with the pcDNA5/FRT/TO vector containing the sequence of ^FLAG^OCIAD2 and the pOG44 vector using GeneJuice (Novagen) as transfection reagent. 24 hours after transfection cells were selected with 200µg/ml HygromycinB (ThermoFisher) to select stable transformants. Individual colonies were further expanded and screened for ^FLAG^OCIAD2 expression under regulation of tetracycline (1µg/ml). OCIAD2-KO cell line was generated using the CRISPR/Cas9 as described (Ran *et al*., 2013). The guide RNAs targeting OCIAD2 gene was designed using software provided by the laboratory of Prof. Feng Zhang (Ran *et al*., 2013). The selected guide was cloned into the pX330 vector using the oligos 5’ CACCGATGGCTTCAGCGTCTGCTCG 3’, 5’ AAACCGAGCAGACGCTGAAGCCATC 3’. HEK293 cells were transfected with pX330 vector together with an EGFP-containing plasmid (pEGFP-N1; Clontech Laboratories, Inc). Single cells were sorted using a FACS Vantage SE machine and the knockouts confirmed by sequencing, SDS–PAGE and Western blotting.

### siRNA mediated knockdown

Cells were reverse-transfected with 25 nM of OCIAD2 siRNA (sense strand: 5’ AGG-UUA-UUU-GGC-AGC-UAA-U55 3’, Eurogentech), 16nM TIMM23 (5’ CCCUCUGUCUCCUUAUUUA 3’, Eurogentech) or 16nM TIM22 (5’ GUGAGGAGCAGAAGAUGAU 3’, Eurogentech) using the Invitrogen Lipofectamine RNAiMAX Reagent (Invitrogen, Carlsbad, CA, USA) diluted with serum-free Gibco Opti-MEM I medium (Gibco, ThermoFisher, Waltham, MA, USA) as per the manufacturer’s recommended protocol. On the third day post-transfection, cells were collected for subsequent assays. As a control, the cells were transfected with Mission siRNA universal negative control (SIC001, Sigma).

### Proliferation assay

24 h prior to the experiment, WT and OCIAD2-KO cells were seeded into six-well format at concentration of 0.1 × 10^6^/well in the growth medium. At the day 0 and then after 24h, 48h and 72h of culture at 37°C with 5% CO_2_ cells were harvested by trypsinization and counted in a cell counter (Countess II, Life Technologies). Cell growth was expressed in absolute cell count numbers.

### Electrophoresis and immunoblot analysis

For Blue Native Page mitochondrial pellets were solubilized in digitonin containing buffer (1% digitonin, 20 mM Tris/HCl, pH 7.4, 0.1 mM EDTA, 50 mM NaCl, 10% (w/v) glycerol, and 1 mM PMSF) to a final concentration of 1µg/µl for 30 min at 4 °C or DDM containing buffer(). Lysates were cleared by centrifugation at 14000x *g* for 10 min at 4 °C followed by adding loading dye (5% Coomassie brilliant blue G-250, 500 mM 6-aminohexanoic acid, and 100 mM Bis-Tris pH 7.0). Samples were loaded onto self-made 4–13% or 4-10% polyacrylamide gradient gels separated and transferred onto PFDV membranes (Merck Millipore) followed by detection by specific antibodies. The High Molecular Weight Calibration Kit for native electrophoresis (Amersham) was used as a molecular weight standard.

### Mitochondria isolation and mitochondrial procedures

Cells were harvested in PBS and suspended in THE buffer (300 mM Trehalose; 10 mM KCl; 10 mM HEPES, pH 7.4; 1 mM EDTA; 0.5 mM PMSF) with 0.1 mg BSA/ml. Cells were homogenized twice using a Potter S dounce homogenizer (Sartorius). After each homogenization step cells were pelleted at 400× *g*, 10 min at 4°C. The supernatant was collected and remaining debris removed by centrifugation (10 min, 4°C, 800× *g*). The supernatant was collected and mitochondria were recovered by centrifugation at 11.000x*g* for 10 min, then washed with BSA-free THE and finally suspended in THE BSA-free buffer. Protein determination was determined by Bradford. The mitochondrial pellet was suspended in BSA-free TH buffer or stored at −80°C until further use. Membrane integration of proteins was determined by 30 min incubation of mitochondria with freshly prepared Na_2_CO_3,_ pH=10.8, followed by centrifugation for 30 min at 100 000x *g*, 4 °C. Submitochondrial localization was analyzed by a protease protection assay. Mitochondrial pellets were resuspended in in either sucrose buffer (250 mM sucrose and 20 mM HEPES/KOH [pH 7.4]) or swelling buffer (25 mM sucrose and 20 mM HEPES/KOH [pH 7.4). Samples were split and either left untreated or treated with proteinase K (50 µg/ml) for 5 min on ice. Reaction was stopped by the addition of 2 mM PMSF.

### Antibodies

The antibodies against following proteins were used in the study: ALR (Santa Cruz Biotechnology, Sc-134869, 1:500), ATP5A (Abcam, ab14748, 1:500), ATP5B (Rabbit serum, Rehling laboratory, 1:500), COA7 (Sigma, HPA029926, 1:500), COX1 (Rabbit serum, Rehling laboratory, 1:2000), COX4 (Cell Signaling Technology, 4850, 1:2000), Cytochrome B (Rabbit serum, Rehling laboratory, 1:1000), (HSP60 (Sigma, H4149, 1:500), MIC60 (Novus Biologicals, NB100-1919, 1:1000), MRPL1 (Rabbit serum, Rehling laboratory, 1:1000), MRPL55 (Proteintech, 17679-1-AP, 1:500), MRPS18(Proteintech 16139-1-AP, 1:500), ND1 (Rabbit serum, Rehling laboratory, 1:1000), ND2 (Rabbit serum, Rehling laboratory, 1:1000), NDUFS1 (Santa Cruz Biotechnology, Sc-50132, 1:1000), Prohibitin 2 (Sigma Aldrich, HPA039874, 1:1000), OCIAD2 (Rabbit serum, Chacińska laboratory, 1:1000), Prohibitin 2 (Sigma Rieske (Rabbit serum, Rehling laboratory, 1:1000), TIMM22 (Proteintech, 14927-1-AP, 1:500), TIMM23 (BD Biosciences, 611222, 1:1000), TIMM29 (Proteintech, 25652-1-AP, 1:500), TOMM20 (Santa Cruz Biotechnology, Sc-11415, 1:500), Tubulin (Santa Cruz Biotechnologys, sc-134239, 1:2000), SDHA (Santa Cruz Biotechnology, Sc-166947, 1:1,000), UQCC2 (Novus Biological, NBP2-14240, 1:500), UQCRB (Sigma Aldrich, HPA002815, 1:500),UQCR1 (Sigma, HPA002815, 1:500).

### Immunopurification of proteins

Isolated mitochondria were solubilized in digitonin containing buffer (150 mM NaCl, 10% glycerol (v/v), 20 mM MgCl_2_, 1 mM PMSF, 50 mM Tris–HCl, pH 7.4, with 1% digitonin (v/w)) at a protein concentration of 1 μg/μl on a rotary wheel for 30–60 min at 4°C. Mitochondrial lysates were cleared by centrifugation at 20,000× *g* at 4°C and transferred to pre-equilibrated anti-FLAG M2 agarose beads (Sigma-Aldrich) for FLAG immunoprecipitation or onto protein A-Sepharose (PAS) containing crosslinked OCIAD2 antibody in a Mobicol spin column (MoBiTec) and incubated at 4°C for 2h under mild agitation. The unbound proteins were removed by spinning out the supernatant for 1 min, 150 × *g* at 4°C. The beads were washed eight times with washing buffer (150 mM NaCl, 10% glycerol (v/v), 20 mM MgCl_2_, 1 mM PMSF, 50 mM Tris–HCl, pH 7.4, with 0.1% digitonin (v/v)). The column-bound proteins were eluted with either 3X FLAG peptide for mass spectrometry or with 0.2 M glycine, pH 2.5 for Western blotting.

### Immunofluorescence

For immunofluorescence, cells were fixed with prewarmed (37°C) 4% paraformaldehyde (Sigma-Aldrich) for 10 min and permeabilised by incubation with PBS containing 0.5% Triton X-100 and blocked with 5% BSA in PBS for 20 min. Subsequently, the cells were incubated with affinity purified anti-OCIAD2 serum or monoclonal mouse anti-FLAG or cyclophilinD. Following washing cells were incubated with Alexa fluor 488 or 568 conjugated secondary antibodies. For mitochondria staining cells were incubated with Mitotracker™ Orange (500nM) for 15 min at 37°C prior fixation and following incubation with secondary antibodies. Confocal images were recorded with a Leica SP8 confocal microscope (Leica Microsystems, Wetzlar, Germany). Z-image series were taken and maximum projections of the stacks are displayed.

### Radioactive precursor synthesis and *in organello* import

Radiolabeled OCIAD2, TIMM22, TIMM23, UQCRB, NDUF13 precursors were synthesized using rabbit reticulocyte lysate (Promega) in the presence of [^35^S] methionine. Isolated mitochondria were diluted in import buffer [250 mM sucrose, 80 mM postassium acetate, 5 mM magnesium acetate, 5 mM methionine, 10 mM sodium succinate, 20 mM Hepes/KOH (pH 7.4), supplemented with 2 mM ATP]. Import reactions were initiated by the addition either of 2% lysate (for SDS-Page analysis) or 8% lysate (assembly on BN-Page) and samples were incubated for the indicated times. To stop the reaction, membrane potential was dissipated on ice using 8 mM antimycin A, 1 mM valinomycin and 10 mM oligomycin. Non-imported proteins were removed by proteinase K (20 μg/mL) treatment for 10 min on ice. PMSF (2 mM) was added to inactivate proteinase K for 10 min on ice. Mitochondria were collected, washed with SEM buffer [250 mM sucrose, 1 mM EDTA, 20 mM Mops (pH 7.2)] and used for SDS-PAGE or BN-Page analyses. Results were visualized using digital autoradiography.

### High-resolution Respirometry and OXPHOS complexes activities

Oxygen consumption was measured in intact or permeabilized HEK293 WT or OCIAD2-KO cell line using Oxygraph-2 k high-resolution respirometer (Oroboros Instruments, Innsbruck, Austria). Data were digitally recorded using DatLab version 5.1.0.20 (Oroboros Instruments, Innsbruck, Austria) expressed as pmol of O_2_/min/10^6^ cells and then normalized to protein content. Before each assay air calibration and background correction were done according to the manufacturer’s protocol. For endogenous cell respiration intact cells were trypsinised, suspended at 0.5 ×10^6^ cells/ml in 2 ml in MiR05 medium (EGTA 0.5 mM, MgCl_2_ 3mM, lactobionic acid 60 mM, taurine 20 mM, KH_2_PO_4_ 10 mM, HEPES 20 mM, D-sucrose 110 mM, 1 mg/ml BSA-fatty acid free added freshly, pH 7.1). Cell suspension was immediately placed into the Oxygraph chamber to measure endogenous respiration. Respiration connected to proton leak was determined by adding ATP synthase inhibitor Oligomycin (final concentration 1.25μM). Subsequently, the mitochondrial inner membrane uncoupler CCCP (Carbonyl cyanide m-chlorophenyl hydrazone) was titrated (0.20 μM/step) to reach maximum respiration (electron transport system capacity, ETS capacity). The complex I inhibitor rotenone (0.5μM) and the complex III inhibitor antimycin A (5μM) were subsequently added to detect oxygen consumption related to each complex and to completely block mitochondrial respiration, showing residual oxygen consumption (ROX). Stable plateaus of oxygen flux in each experimental step were corrected for ROX afterwards.

Substrate-driven respiration was assessed in digitonin-permeabilized cells. CI driven respiration and ATP synthesis, were assessed by adding 5mM pyruvate in combination with 0.5mM malate, as respiratory substrates. To detect complex III linked to OXPHOS activity 5 mM Glycerol-3-phosphate was added prior Malate and Pyruvate. Cytochrome c (10 μM) was added in parallel experiments to test for the intactness of the mitochondrial outer membrane in digitonin-treated cells. Complex IV activity was measured as previously described (Mohanraj *et al*., 2019). In brief: digitonized cellular extracts from the OCIAD2 -KO and WT cells were added to 200 μl of 100 μM reduced cytochrome *c* in 50 mM potassium phosphate buffer (pH 7.0) and solution was pipetted into 96-well format plate. Decrease in absorbance at 550 nm was recorded for 3 min, which corresponded to linear changes in absorbance. Specific activity of complex IV was calculated by subtracting the absorbance change in the blank sample. The protein concentration of digitonized cell suspension was measured by Bradford assay, and complex IV activity was expressed in nanomoles of cytochrome *c* per minute per milligram of protein. Data were analyzed by Student’s t-test. Data are presented as means ± SD of 3 experiments. *p = 0.05, ** = p < 0.01, *** = p < 0.001.

### Preparation of samples for proteomic analyses

Samples for LFQ analysis were prepared according to Sample Preparation by Easy Extraction and Digestion (SPEED) protocol (Doellinger *et al*., 2020). In brief, 50 µg of mitochondrial pellets were resuspended in trifluoroacetic acid (TFA) (sample/TFA 1:4 (v/v) and incubated at RT for 5 min. Samples were then neutralized with 2M TrisBase using 10 x volume of TFA and further incubated at 95°C for 10 min after adding Tris(2-carboxyethyl)phosphine (TCEP) to the final concentration of 10 mM and 2-chloroacetamide (CAA) to the final concentration of 40 mM. Samples were then diluted with water (1:5) and incubated overnight at 37°C with 0.5 µg of sequencing grade modified trypsin (Promega). Samples were acidified using 1% TFA and peptides separated into 3 fractions on a 200 µL StageTips packed with three Empore™ SPE Disks SDB-RPS (3M) according to a previously published protocol (Kulak *et al*., 2014), and concentrated using a Savant SpeedVac concentrator. Prior to LC-MS measurement, the samples were resuspended in 0.1 % TFA, 2% acetonitryl in water.

For SILAC analysis, affinity enriches proteins eluted with FLAG peptide (final concentration 1 mg/mL) were diluted in 0.1 M Tris pH 8.0 containing 5mM TCEP and 10mM 2-chloroacetamide, and incubated overnight at 37°C with 0.5 µg of sequencing grade modified trypsin (Promega). Samples were acidified using 1% TFA and desalted using 200 µL StageTips packed with three Empore™ SPE Disks C18 (3M) (Rappsilber *et al*., 2007) and concentrated using a Savant SpeedVac concentrator. Prior to LC-MS measurement, the samples were resuspended in 0.1 % TFA, 2% acetonitryl in water.

### LC-MS/MS analysis

Chromatographic separation was performed on an Easy-Spray Acclaim PepMap column 25cm or 50 cm long × 75 µm inner diameter (Thermo Fisher Scientific) at 35 °C by applying a 60 min (mitochondrial protein levels) or a 90 min (Flag-OCIAD2 co-IP) acetonitrile gradients in 0.1% aqueous formic acid at a flow rate of 450 or 300 nl/min, respectively. An UltiMate 3000 nano-LC system was coupled to a Q Exactive HF-X mass spectrometer via an easy-spray source (all Thermo Fisher Scientific). For the Flag-OCIAD2 co-IP experiments, the Q Exactive HF-X was operated in data-dependent mode with survey scans acquired at a resolution of 60,000 at m/z 200. Up to 12 of the most abundant isotope patterns with charge 2 or higher from the survey scan were selected with an isolation window of 1.3 m/z and fragmented by higher-energy collision dissociation (HCD) with normalized collision energies of 27, while the dynamic exclusion was set to 15 s. The maximum ion injection times for the survey scan and the MS/MS scans (acquired with a resolution of 15,000 at m/z 200) were 45 and 22 ms, respectively. The ion target value for MS was set to 3×10e6 and for MS/MS to 10e5, and the intensity threshold for MS/MS was set to 8.3×10e2. For the determination of the mitochondrial protein levels, the Q Exactive HF-X was operated in a ‘BoxCar’ mode through MaxQuant Live (Wichmann *et al*., 2019) with survey scans acquired at a resolution of 120,000 at m/z 200 within m/z range 350-1650, followed by 3 BoxCarScans (12 BoxCarBoxes) within m/z range 400-1200. Up to 5 of the most abundant isotope patterns with charge states 2-5 from the survey scan were selected with an isolation window of 1.4 m/z and fragmented by higher-energy collision dissociation (HCD) with normalized collision energies of 27, while the dynamic exclusion was set to 40 s. The maximum ion injection times for the survey scan and the MS/MS scans (acquired with a resolution of 15,000 at m/z 200) were 250and 28 ms, respectively. The ion target value for MS was set to 10e6 and for MS/MS to 10e5.

### Proteomics data processing

The data were processed with MaxQuant v. 1.6.6.0 (Cox and Mann, 2008), and the peptides were identified from the MS/MS spectra searched against Human Reference Proteome UP000005640 using the build-in Andromeda search engine. Cysteine carbamidomethylation was set as a fixed modification and methionine oxidation as well as protein N-terminal acetylation were set as variable modifications. For in silico digests of the reference proteome, cleavages of arginine or lysine followed by any amino acid were allowed (trypsin/P), and up to two missed cleavages were allowed. The FDR was set to 0.01 for peptides, proteins and sites. For SILAC samples (Flag-OCIAD2 Co-IP samples), multiplicity of 2 was selected and light labels set as Arg0 & Lys0, and heavy labels set as Arg10 & Lys8, the maximal number of labelled amino acids per peptide set to 3, and requantify option was set to on. Match between runs was enabled. Other parameters were used as pre-set in the software. Unique and razor peptides were used for quantification enabling protein grouping (razor peptides are the peptides uniquely assigned to protein groups and not to individual proteins). Data were further analysed using Perseus version 1.6.6.0 and Microsoft Office Excel 2016.

### LFQ-based differential analysis of mitochondrial protein levels

LFQ values for protein groups were loaded into Perseus v. 1.6.6.0. Standard filtering steps were applied to clean up the dataset: reverse (matched to decoy database), only identified by site, and potential contaminant (from a list of commonly occurring contaminants included in MaxQuant) protein groups were removed. Protein accessions were matched to accessions generated from gene names constituting the IMPI data set (Smith and Robinson, 2019), and successfully matched protein groups retained for further analysis. LFQ intensities were log2 transformed, protein groups with LFQ values in less than 3 samples were removed, and all remaining missing values were imputed from normal distribution (width = 0.3, down shift = 1.8 × standard deviation). Gaussian distribution of log2 transformed LFQ intensities were confirmed by histogram analysis preventing the unbiased introduction of small values. Student’s T-test (permutation-based FDR with 250 randomizations = 0.01, S0 = 0.25) was performed to return proteins which levels were statistically significantly changed in response to OCIAD2-KO (Table S2).

### SILAC-based Flag-OCIAD2 Co-IP enrichment analysis

Ratio H/L normalized values for protein groups were loaded into Perseus v. 1.6.6.0. Standard filtering steps were applied to clean up the dataset: reverse (matched to decoy database), only identified by site, and potential contaminant (from a list of commonly occurring contaminants included in MaxQuant) protein groups were removed. Protein accessions were matched to accessions generated from gene names constituting the IMPI data set (Smith and Robinson, 2019), and successfully matched protein groups retained for further analysis. Ratio H/L normalized values were log2 transformed, and mean fold enrichment values for Flag-OCIAD2 vs Ctrl were calculated for all protein groups; the mean fold enrichment threshold for was set to 2 (Table S1).

## General

We thank Sven Dennerlein, Urszula Nowicka and Michał Wasilewski for scientific discussion and Albert Roethel for experimental assistance.

## Supplementary Tables

**Table S1:** Silac analysis of FLAGOCIAD2.

**Table S2:** LFQ isolated mitochondria.

## FIGURE AND LEGENDS

**Supplementary Figure 1:**
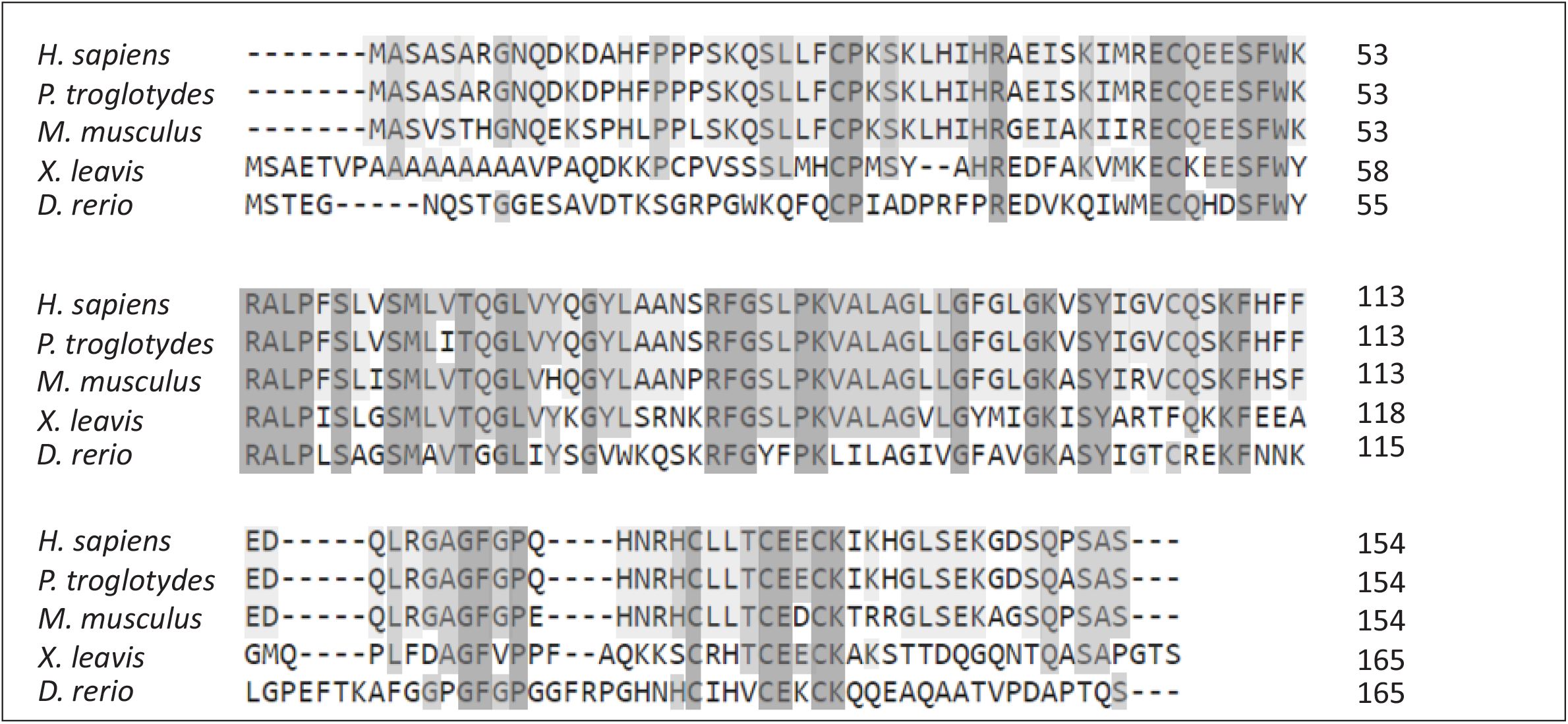
OCIAD2 is a metazoan specific protein. Alignment of the human (H. sapiens) OCIAD2 amino acid sequence to its metazoan homologs using ClustalW. Dark gray shading indicates identical amino acids, whereas medium or light gray shading depicts similar amino acids: 80% or 60% similarity, respectively.

**Supplementary Figure 2:**
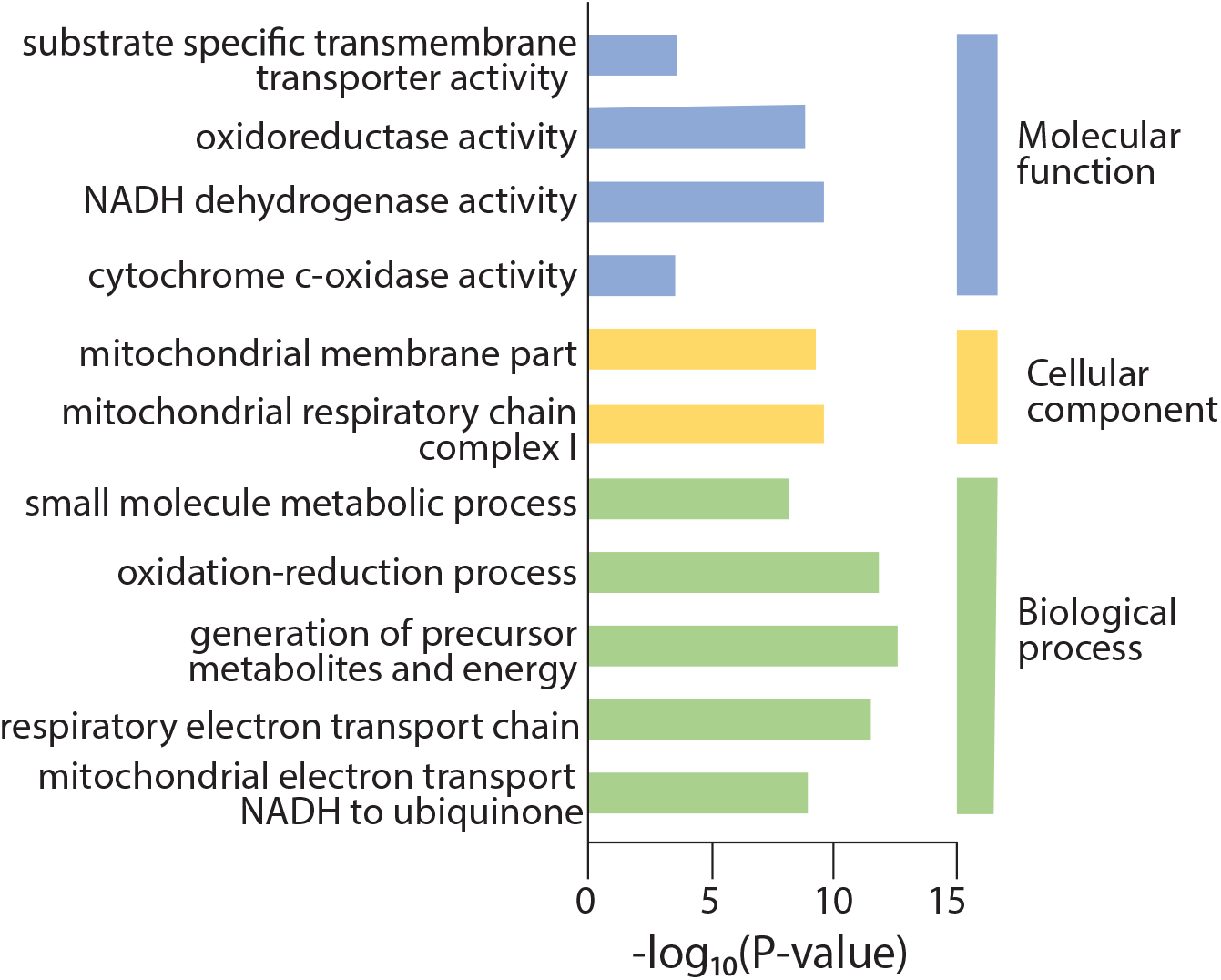
OCIAD2 interacts with electron transport chain proteins. Go terms significantly enriched among the mitochondrial proteins specifically bound to ^FLAG^OCIAD2 obtained in Fisher test’s for 48 proteins groups with mean fold enrichment >2 in ^FLAG^OCIAD2 vs Ctrl samples compared to all 180 IMPI listed protein groups (bait was escluded) identified in the co-IP experiment (FDR=0.01).

